# Identification and characterization of tertiary lymphoid structures in brain metastases

**DOI:** 10.1101/2024.09.30.614674

**Authors:** Sadaf S. Mughal, Yvonne Reiss, Jörg Felsberg, Lasse Meyer, Jadranka Macas, Silja Schlue, Tatjana Starzetz, Karl Köhrer, Tanja Fehm, Volkmar Müller, Katrin Lamszus, Dirk Schadendorf, Iris Helfrich, Harriet Wikman, Anna Berghoff, Benedikt Brors, Karl H. Plate, Guido Reifenberger

**Affiliations:** Division Applied Bioinformatics, German Cancer Research Center (DKFZ), Im Neuenheimer Feld 280, 69120 Heidelberg, Germany; Institute of Neurology (Edinger-Institute), University Hospital, Goethe University, Heinrich-Hoffmann-Strasse 7 60590 Frankfurt, Germany; Frankfurt Cancer Institute (FCI), Paul-Ehrlich-Straße 42-44, 60596 Frankfurt, Germany; Institute of Neuropathology, Medical Faculty, Heinrich Heine University and University Hospital Düsseldorf, Moorenstrasse 5, Düsseldorf, Germany; Center for Biological and Medical Research (BMFZ), Genomics and Transcriptomics Laboratory (GTL), Heinrich Heine University, Universitätsstrasse 1, Düsseldorf, Germany; Department of Gynecology and Obstetrics, Medical Faculty, Heinrich Heine University and University Hospital Düsseldorf, Moorenstrasse 5, Düsseldorf, Germany & Center of integrated Oncology ABCD.; Department of Gynecology, University Medical Center Hamburg-Eppendorf, Martinistrasse 52, 20246 Hamburg, Germany; Laboratory for Brain Tumor Biology, Department of Neurosurgery, University Medical Center Hamburg-Eppendorf, Martinistrasse 52, 20246 Hamburg, Germany; Department of Dermatology, University Hospital Essen, University Duisburg-Essen, Hufelandstrasse 50, 45147 Essen, Germany; Department of Dermatology and Allergy, University Hospital of Munich, Ludwig-Maximilian-University (LMU), Frauenlobstrasse 9-11, 80337 Munich, Germany; Department of Tumor Biology, University Medical Center Hamburg-Eppendorf, Martinistrasse 52, 20246 Hamburg, Germany; Department of Internal Medicine 1, Clinical Division of Oncology, Medical University Vienna, Währinger Gürtel 18-20, 1090 Vienna, Austria; Medical Faculty and Faculty of Biosciences, Heidelberg University, Germany; Faculty of Biosciences, Heidelberg University, 69120 Heidelberg, Germany; National Center for Tumor Diseases (NCT), Im Neuenheimer Feld 410, 69120 Heidelberg, Germany; German Cancer Consortium (DKTK), Core Center Heidelberg, Germany; German Cancer Consortium (DKTK), Partner site Frankfurt/Mainz and German Cancer Research Center (DKFZ), Heidelberg, Germany; German Cancer Consortium (DKTK), Partner Site Essen/Düsseldorf and German Cancer Research Center (DKFZ), Heidelberg, Germany; German Cancer Consortium (DKTK), Partner Site Munich and German Cancer Research Center (DKFZ), Heidelberg, Germany

**Keywords:** Brain metastasis, immune cell deconvolution, multiplex immunofluorescence, prognosis, tertiary lymphoid structures, tumor microenvironment

## Abstract

Brain metastases (BrM) are the most common cancers in the brain. We performed transcriptome-wide gene expression profiling combined with spatial immune cell profiling to characterize the tumor immune microenvironment in BrM from different primary tumors. We found that BrM from lung carcinoma and malignant melanoma showed overall higher immune cell infiltration as compared to BrM from breast carcinoma. RNA sequencing-based immune cell deconvolution revealed gene expression signatures indicative of tertiary lymphoid structures (TLS) in subsets of BrM, mostly from lung cancer and melanoma. This finding was corroborated by multiplex immunofluorescence staining of immune cells in BrM tissue sections. Detection of TLS signatures was more common in treatment-naïve BrM and associated with prolonged survival after BrM diagnosis in lung cancer patients. Our findings highlight the cellular diversity of the tumor immune microenvironment in BrM of different cancer types and suggest a role of TLS formation for BrM patient outcome.

## Introduction

Hematogenous metastasis formation is a multistep process that involves cancer cell invasion into the vascular system, dissemination via the blood stream, as well as endothelial adhesion and vascular extravasation, followed by invasive growth in the distant organ^1^. The brain is a frequent site of hematogenous metastases from various types of cancers as indicated by autopsy studies reporting the presence of brain metastases (BrM) in approximately 25 % of patients who died of cancer^2^. In total, BrM are approximately 10-times more common than primary brain tumors, making them the overall most frequent central nervous system tumors in adults^3,4^.The most common primary cancers giving rise to BrM are lung cancer (20-56%), breast cancer (5-20%) and melanoma (7-16%), followed by renal cell carcinoma (RCC) and colorectal cancer^3,5,6,7^. Despite remarkable therapeutic advances, including novel molecularly targeted approaches and immune checkpoint inhibition, metastases still constitute the primary cause of cancer-related mortality^8^. In line, clinical outcome of the vast majority of patients with BrM remains dismal, with reported median overall survival rates of 8.1% and 2.4% after 2 and 5 years, respectively^9,10^. This underscores the urgent need to gain a deeper understanding of the molecular and cellular pathomechanisms in BrM to devise more effective treatment strategies.

In the present study, we aimed to further decipher the complex tumor immune microenvironment in BrM of different primaries, with a specific focus on the characterization of diverse immune cell populations orchestrating the formation of specialized lymphoid architectures resembling tertiary lymphoid structures (TLS) within these tumors. TLS are ectopically formed aggregates of lymphoid and stromal cells, comprising T cell zones with antigen-presenting dendritic cells, and B cell zones with germinal centers in different stages of maturity^11,12^. Formation of TLS or lymphoid aggregates has been detected in various cancer types and has been implicated as an independent predictor of immunotherapy response and prognosis across several tumor types including melanoma^13,14,15^, non-small cell lung cancer (NSCLC)^16^, RCC^13^ and sarcomas^17^. Despite these recent insights into the important roles of TLS in various cancer types, their presence, functional roles, and clinical significance in BrM patients have remained elusive to date. We therefore performed comprehensive molecular and cellular analyses of tumor tissues of 95 patients with BrM originating from carcinomas of the lung, breast, kidney and colon as well as from cutaneous melanoma. Utilizing bulk RNA sequencing-based immune cell deconvolution, we observed variable levels of tumor-infiltrating leukocytes and the presence of TLS-like gene expression signatures in approximately one-third of the analyzed BrM tissues. These signatures were most frequently observed in BrM derived from lung cancer and melanoma, whereas BrM from breast cancer exhibited comparatively lower expression of TLS signatures. Using multiplex immunofluorescence (mIF) combined with spatial imaging we confirmed the presence of TLS or TLS-lymphoid aggregates in subsets of BrM at the cellular level. In addition, we revealed that gene expression signatures indicative of TLS formation were more frequently detected in treatment-naïve BrM tumor tissues and associated with prolonged survival of lung cancer patients following BrM diagnosis.

## Results

### Detection of heterogeneous populations of tumor-infiltrating immune cells in brain metastases

We performed whole transcriptome sequencing on RNA extracted from BrM tissue samples of 95 patients (Table S1, Fig. S1) and applied reference-based deconvolution using MCP-counter^18^ to assess the immune cell infiltrates in each tumor (Table. S2). Unsupervised clustering using immune cell-based deconvolution assigned the BrM tissues to one of six MCP-counter-based immune classes, labelled IC1 to IC6 (Fig. 1A). The pattern of immune cell infiltration was markedly distinct for the six IC classes, with IC1 tumors showing the overall lowest infiltration by immune and stromal cells, while the other IC classes demonstrated differently composed immune cell infiltrates with variable contributions of the distinct lymphoid, myeloid and stroma cell types (Fig. 1A). We observed varying degrees of immune cell abundance according to the primary origin of BrM, with lung cancer BrM showing the overall highest ICI scores, while breast cancer BrM demonstrating the overall lowest ICI scores that were significantly lower when compared to melanoma and lung BrM (Fig. 1B). Interestingly, compared to IC1 class tumors, tumors assigned to IC2-IC6 classes showed significantly higher abundance scores for T cells, B cells, and myeloid dendritic cells, all crucial components of TLS formation (Fig.1A. Fig. S2A).

**Figure 1.**
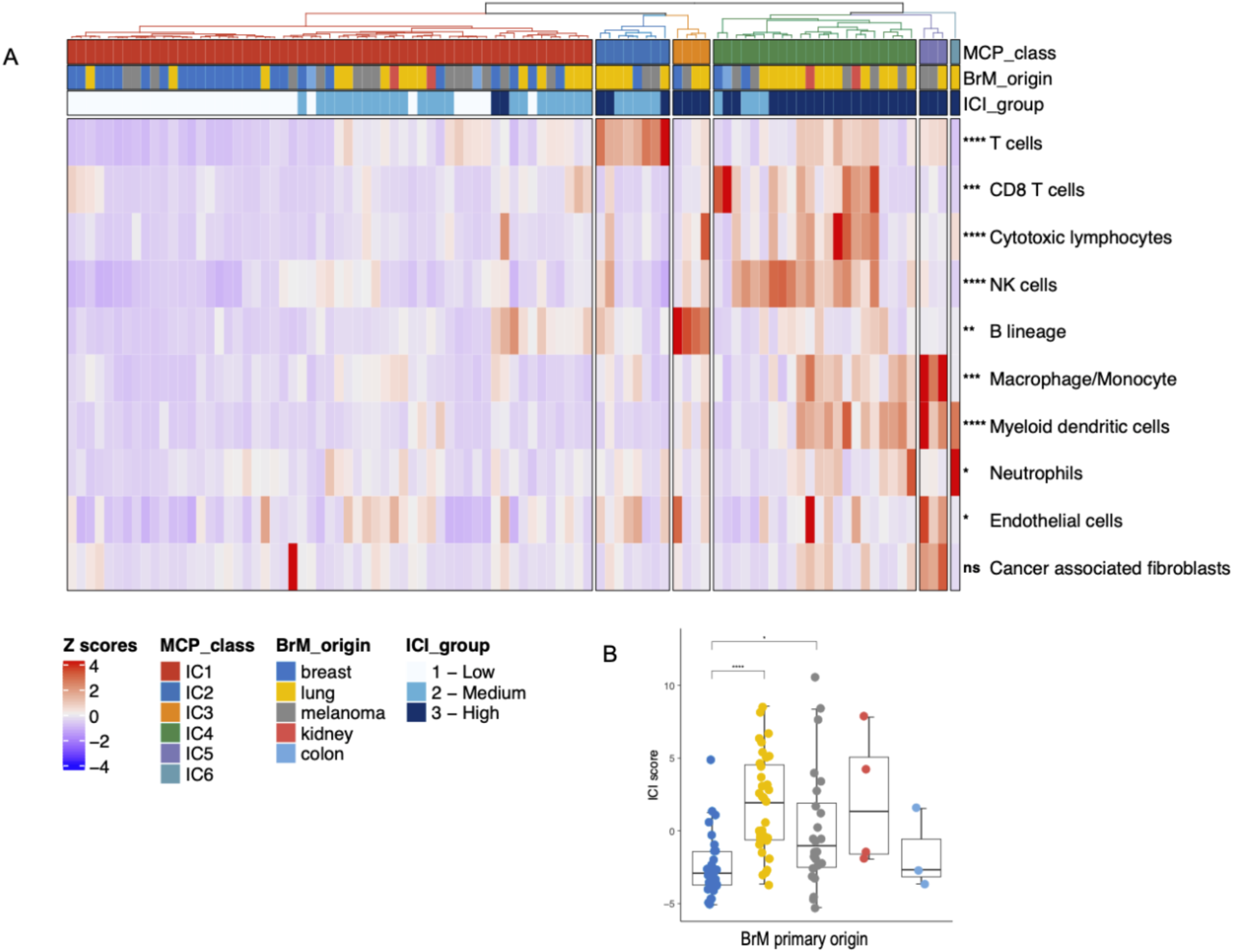
Bulk RNA sequencing-based immune cell deconvolution reveals heterogeneous cell populations in brain metastases of different primary origins. **(A)** Heatmap displaying the immune and stromal composition of brain metastases. Unsupervised hierarchical clustering of samples based on MCP-counter scores for the respective immune cell types shows six distinct immune classes (ICs). NK cells, Natural killer cells. (B) Comparison of Immune cell infiltration (ICI) scores among brain metastases from different origins (breast, lung, melanoma, kidney, colon). ICI scores were calculated based on the sum of MCP scores for immune cells only, excluding endothelial cells and fibroblast populations. Samples were assigned into high, intermediate and low immune cell infiltration groups based on tertiles scores. Statistical testing was performed using a two-sided Kruskal-Wallis test. Significance levels between groups are: ****, p≤0.0001; ***, p≤0.001; **, p≤0.01; *, p≤0.05; ns (not significant), p>0.05. The p-values are corrected for multiple testing using Benjamini-Hochberg correction. Row and column clustering were enabled using the agglomerative hierarchical clustering method Agnes with an Euclidian distance metric and Wards linkage criterion. A legend illustrating the color coding of samples is provided on the left bottom.

### Identification of gene signatures of tertiary lymphoid structures (TLS) in BrM with immune cell-rich microenvironment

We constructed a 32-gene signature by integrating components from the TLS hallmark^14^, the 12 chemokine signature^19^, and surrogate markers for Th1 cells, Tfh cells, B cells and dendritic cells to reveal transcriptomic evidence of TLS formation in BrM tissues (Fig. S2B and Table S3). Unsupervised hierarchical clustering based on TLS signature genes classified the investigated BrM samples into three distinct clusters: TLS1 – “*low expression*”, TLS2 – *“intermediate expression”,* TLS3 – “*high expression*” (Fig. 2A). The majority of breast cancer BrMs were assigned to the TLS1 cluster (68%), with only a single sample assigned to the TLS3 group. Melanoma and lung cancer BrMs exhibited significant heterogeneity, with samples distributed across all three TLS clusters. However, the TLS3 cluster primarily comprised samples from lung BrM exhibiting the highest TLS signature score (Fig. 2B). In total, the TLS3 cluster included 12/35 BrM from lung cancers (all corresponding to non-small cell lung cancer (NSCLC)), 7/24 BrM from melanoma, 2/4 BrM from renal cell carcinoma, and 1/27 BrM from breast cancer. From the 7 small cell lung cancer (SCLC) BrM, four samples were assigned to the TLS1 cluster and the remaining three to the TLS2 cluster. Cluster assignment of lung cancer BrM did not significantly differ between BrM from SCLC or NSCLC (Fischer’s exact test *P value* = 0.1452), although none of BrM from SCLC were assigned to the TLS3 cluster. The cytolytic activity score was highest among the samples assigned to the TLS3 cluster suggesting a cooperative interaction of cytolytic T cells at TLS sites in the tumor microenvironment (Fig. 2C).

**Figure 2.**
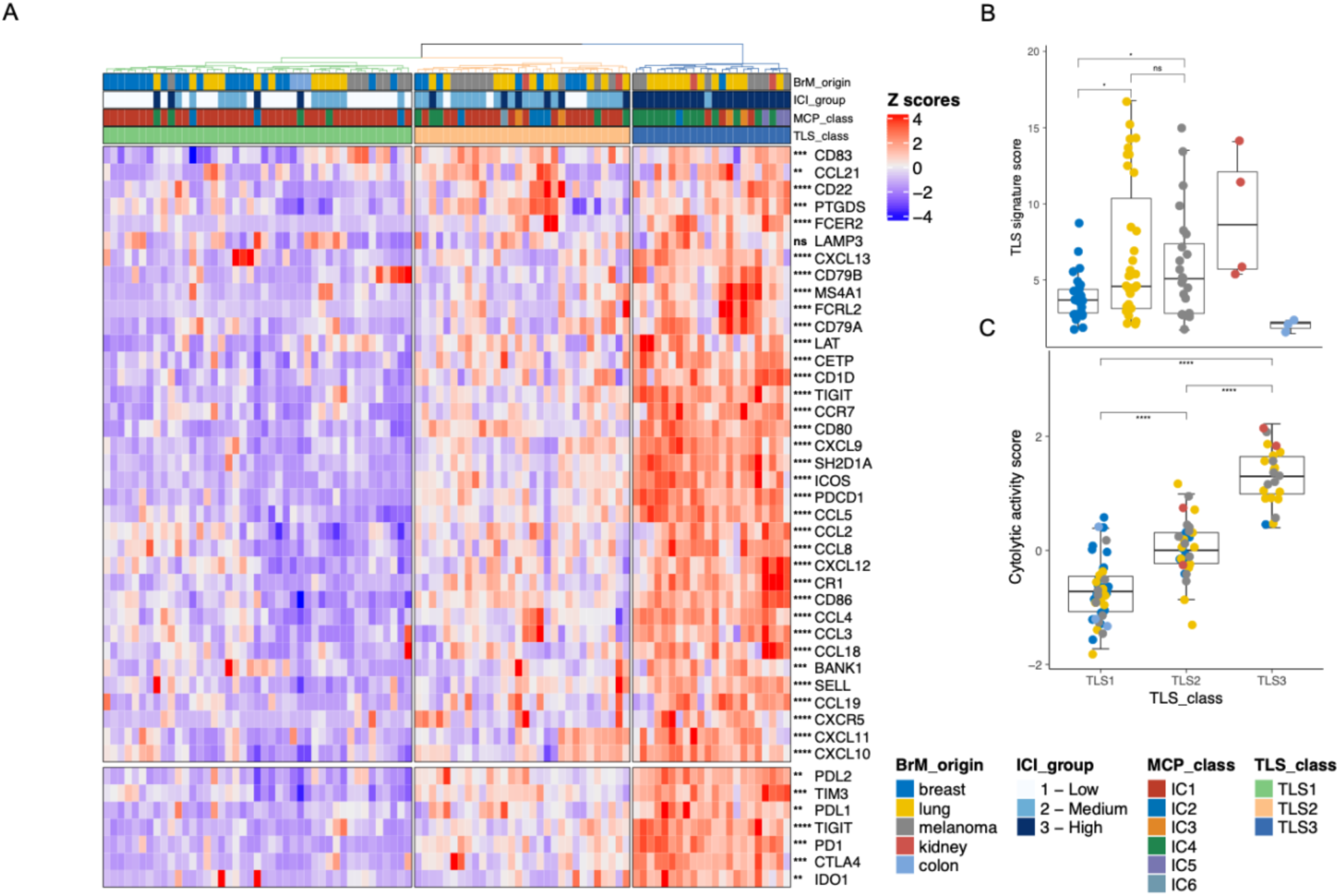
TLS signatures are a feature of the immune-rich BrM tumor microenvironment. **(A)** Heatmap based on unsupervised hierarchical clustering of genes belonging to the TLS signature score. Patients are assigned to one of three TLS classes based on clustering into high (TLS3), intermediate (TLS2) and low expression (TLS1) of TLS signature genes. The bottom panel of the heatmap shows the expression of immunotherapy-relevant markers. Statistical testing was performed using two-sided Kruskal-Wallis tests. Row and column clustering were enabled using the agglomerative hierarchical clustering method Agnes with an Euclidian distance metric and Wards linkage criterion. Legend for color coding is provided (bottom left). **(B)** TLS signature score distribution in BrM according to primary tumor type. The TLS score was derived by calculating the geometric means of the TLS signature genes. The boxplots display the distribution of TLS scores among BrM of different primary tumors. **(C)** Cytolytic score distribution in BrM according to the TLS classes. Cytolytic score is the log-average of *GZMA* and *PRF1* normalized gene expression. Tumors belonging to the TLS3 (TLS high) class showed an overall higher cytolytic activity compared to TLS2 and TLS1 classes. Statistical testing was performed using an unpaired two-sided Wilcoxon test. The p-values are corrected for multiple testing using Benjamini-Hochberg correction. Legend for color coding is provided. Significance levels between groups are: ****, p≤0.0001; ***, p≤0.001; **\*\*,** p<0.01; *, p≤0.05; ns (not significant), p>0.05.

### Detection of TLS in BrM by spatial proteome profiling

To further investigate the cellular immune landscape and validate the presence of TLS as identified through transcriptomic analyses, we performed multispectral immunofluorescence (mIF) imaging and spatial analyses on 60 samples of our BrM cohort, selected based on the availability of adequate tissue specimens. We established a 7-plex mIF panel to detect lymphoid aggregates and TLS using markers against T cells (CD3), B cells (CD20), mature dendritic cells (Lamp-3), macrophages (CD163), as well as endothelial cells (vWF) and tumor cells (pan-cytokeratin or MelA) (Fig. 3A). Frequencies and distribution patterns of B and T lymphocytes across BrM from cases of lung, breast, kidney, colon carcinoma and cutaneous melanoma are displayed in the heat map shown in Fig. 3B. High numbers of B cells and TLS-like lymphoid aggregates were most prominent in subsets lung cancer BrM samples (Fig. 3B, Fig. S3), thus confirming the bulk RNA sequencing results. Multispectral images in Fig. 3C show examples of B cell aggregates and TLS scores in different BrM ranging from high to low (evidenced by spatial blots of whole tissue biopsies). Composite and single-plex images of lymphoid aggregates and TLS in selected BrM originating from different primary tumors as well as the frequencies and cellular compositions of these lymphoid structures are displayed in Fig. S3. In an independent patient cohort^20^ comprising 53 patients with melanoma BrM, analysis with the 7-plex TLS panel revealed that 42% of the BrM samples exhibited characteristics indicative of TLS formation (Table S4).

**Figure 3.**
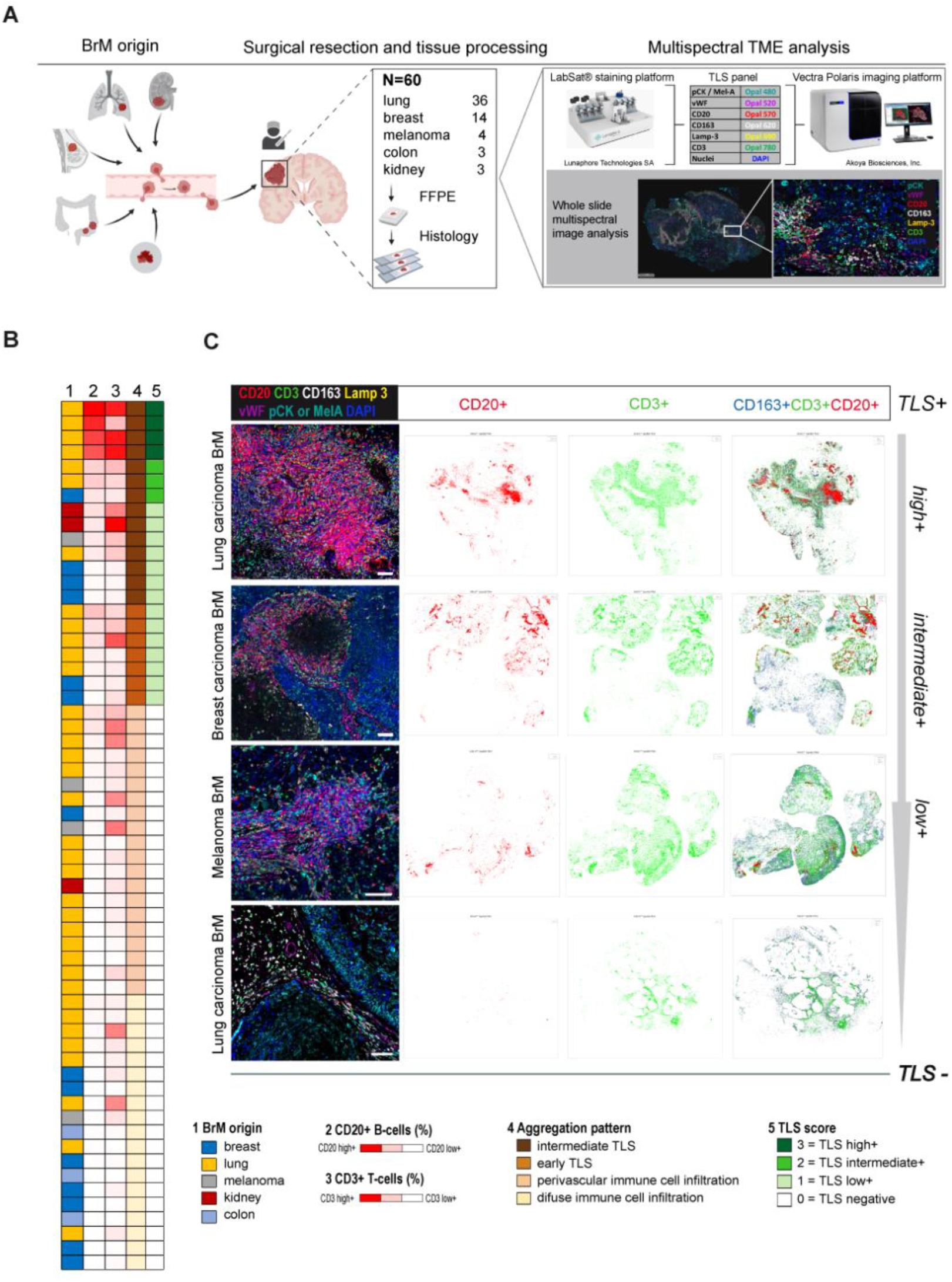
Identification of TLS in BrM tissues by multiplex immunofluorescence and spatial image analysis. **(A)** Cartoon displaying the BrM patient cohort investigated by multiplex immunofluorescence and the multispectral tumor microenvironment (TME) analysis. Formalin-fixed and paraffin-embedded (FFPE) BrM tissue samples of 60 patients were investigated by multiplex immunofluorescence using the antibody panel and technology platform as illustrated (for further details see Materials and Methods). Created with BioRender.com **(B)** Heatmap displaying BrM origins (1), B and T lymphocyte frequencies (2, 3), aggregation pattern (4), and TLS scores (5) in the 60 BrM patient samples. (**C)** Representative composite images showing the Opal multiplex immunofluorescence results employing the 7-plex BrM TLS panel with antibodies directed against CD3, CD20, CD163, Lamp3, vWF, pan-cytokeratin (pCK) or Mel A, and DAPI in selected BrM of different primary tumor origin. Whole slide spatial plots display the distribution of CD20+ and CD3+ leukocytes, and their overlay with CD163+ macrophages in selected cases of a lung carcinoma BrM, a breast carcinoma BrM, and a melanoma BrM with TLS frequencies ranging from high to low. Scale bars: 50 µm.

We found an approximately 55% concordance between the mRNA signature score-based stratification of BrM into distinct TLS classes and the stratification in TLS classes based on multiplex immunofluorescence staining (Fig. S4). In addition, we observed a positive correlation between the MCP-counter-predicted cell scores for B cells, T cells and monocytic lineage cells with high densities of CD20^+^, CD3^+^ and CD163^+^ cells as determined by spatial immune cell profiling (Fig. S4 A and B). Moreover, expression levels of PD-1 and PD-L1 mRNA showed a strong correlation with the abundance of CD3+ T cells and CD20+ B cells in BrM tissue samples (Fig. S5A).

### Characterization of transcriptional hallmarks of TLS-positive BrM

To characterize transcriptional programs in BrM according to the TLS status, we stratified our cohort into TLS-positive and -negative tumors, in which the TLS status was concordant according to gene expression signatures and multiplex immunofluorescence staining. Upon differential gene expression analysis, we found a dominant expression of immunoglobulin (IG) signature genes as well as higher expression of B cell markers like *MS4A1, FCRL2, CD79A* and *CR1* in TLS-positive tumors indicative of mature TLS states (Fig. 4A, Table S5). Several T cell lineage genes (*CD3D, CD28, ICOS, CTLA4*) were also upregulated in TLS-positive tumors, as were cytotoxic T cell markers such as *GZMA, GZMB, GZMH* and *GZMK* indicating robust humoral and cell-mediated immune responses.

**Figure 4.**
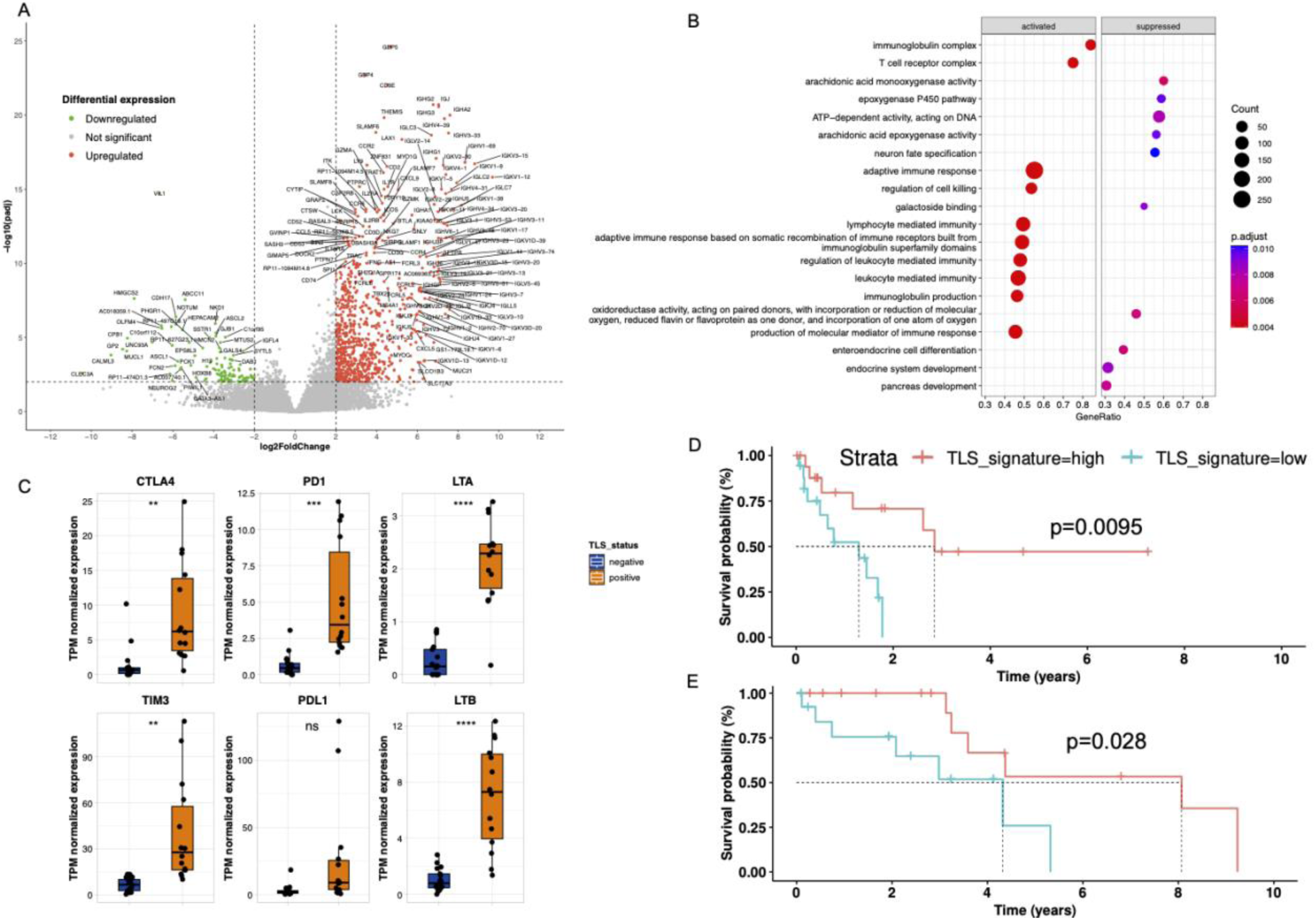
Activated transcriptional programs in the TLS-positive tumors and prognostic impact of TLS in BrM. **(A)** Differentially expressed genes (DEG) between TLS positive and TLS negative tumors using RNAseq data. Volcano plot showing upregulated and downregulated genes in TLS positive tumors (n=31). A cut-off of gene expression fold change of ≥ 2 or ≤ 2 and a false discovery rate (FDR) q ≤ 0.05 was applied to filter DEGs. **(B)** Gene set enrichment analysis based on DEGs displays the activated and suppressed transcriptional programs in TLS positive tumors. The size of the dot represents the gene count and the adjusted p-values are shown as color gradient on the left. **(C)** Expression differences in selected immunotherapy-relevant gene markers and lymphotoxin genes in TLS positive and negative cases. Box plots display TPM normalized gene expression values. Significance levels between groups are: ****, p≤0.0001; ***, p≤0.001; **, p≤0.01; *p≤0.05; ns (not significant), p>0.05. **(D**) Kaplan-Meier survival estimation for lung BrM patients stratified according to RNAseq-based TLS signature score. Overall survival (OS) is calculated from the diagnosis of the resected BrM. TLS status is color-coded on top. The p-value was calculated using the log-rank method. **(E)** Kaplan-Meier survival estimation of OS following the diagnosis of BrM in a published cohort of lung BrM patients^24^. TLS status was determined using the RNA-based TLS signature scores. The p-value was calculated using the log-rank method.

Furthermore, we observed higher mRNA expression levels of the CXCR3 ligand genes *CXCL9* and *CXCL10* in TLS-positive as compared to TLS-negative tumors, suggesting the recruitment of CD8+ T cells into tumor mass^21^. Among the differentially expressed genes, lymphotoxin beta (LTß) expression was significantly upregulated in TLS-positive tumors (Fig. 4C, Fig. S5B), which is of interest as LTB signaling has been linked to the maintenance and functioning of lymphoid homeostasis and certain aspects of TLS development are mediated via LTßR signaling^22,23^.

Gene set enrichment analysis of the differentially expressed transcripts between TLS-positive and - negative BrM revealed activation of adaptive immune cell response, including activation of immunoglobulin and T cell receptor complex, in TLS-positive tumors (Fig. 4B). Similarly, we found an elevated expression of immune checkpoint-related genes (CTLA-4, PD-L1) in TLS-positive tumors hinting towards a potential link between TLS and response to immunotherapy (Fig. 4C).

### Association of TLS with prolonged survival of lung cancer BrM patients

Next, we explored the potential association between the presence of TLS structures and patient outcomes, specifically focusing on patients with lung cancer BrM. This subgroup was the largest within our BrM cohort and composed of reasonable numbers of BrM classified into different TLS classes based on gene expression signatures, facilitating the analysis of this relationship. We stratified our lung BrM patient cohort based on the median expression of the TLS signature score and compared the survival outcomes in lung BrM patients belonging to the TLS high against the TLS low class. Univariate survival analysis in the entire lung cancer BrM patient cohort (n=35 patients) revealed a positive association towards prolonged overall survival (OS) (p=0.0095) following BrM diagnosis in the patients with BrM assigned to the TLS high score (Fig. 4D). In addition, an association with survival was observed in an independently published cohort of patients with lung BrM^24^ that we also stratified according to the tumors’ TLS signature scores. In this cohort, univariate analysis revealed a longer overall survival (log-rank test, p=0.028) from the diagnosis of BrM in patients with TLS high score (Fig. 4E). The positive association with survival was retained in the subgroup of 28 patients with NSCLC BrM from our institutional cohort and the 25 patients with NSCLC BrM in the independently published cohort (Fig. S6A). In addition to the association with patient survival, we also evaluated the potential influence of previous chemotherapy for primary tumor treatment on TLS class assignment of BrM tissues in our institutional cohort. Information on treatment before BrM resection was available from 34/35 lung cancer BrM patients. Among these, 14 patients had been treated before BrM resection while 20 patients had not been treated. Logistic regression using a binominal model showed that the TLS high group was enriched for patients with treatment-naïve BrM (p=0.00835) (Fig. S6B).

## Discussion

The immune-privileged nature of the brain^25^, in conjunction with brain-intrinsic cell types and boundaries formed by the blood-brain and cerebrospinal fluid-brain barriers create an intricate micromilieu that has a profound impact on tumor initiation and growth in the central nervous system as well response to therapy of brain tumors^26,27^. BrM are the most common cancers in the brain and in most patients associated with poor outcome^28^. Therefore, gaining a comprehensive understanding of the molecular and cellular pathomechanisms underlying the development and progression of BrM, particularly the context of the tumor immune microenvironment, is of paramount importance. This knowledge is critical in light of novel therapeutic strategies, such as immune checkpoint inhibition, which have demonstrated clinical efficacy in patients with various cancers, including those prone to brain metastasis, such as melanoma and non-small cell lung cancer^29,30,31^.

We approached characterization of the tumor microenvironment in BrM through deconvolution of bulk RNA sequencing data, which revealed notable differences in immune cell infiltration among BrM originating from distinct types of primary cancers such as lung and breast carcinoma as well as cutaneous melanoma. Specifically, breast BrM demonstrated the lowest degree of immune cell infiltration, while many BrM from lung cancer, in particular BrM from NSCLC, and melanoma BrM exhibited an immune cell-rich microenvironment. These results align with previous findings showing that the tumor immune microenvironment may differ between different types of BrM, with subsets of tumors showing an immune cell-enriched phenotype^32,33^.

Recent studies have demonstrated a crucial role for B cells and TLS with respect to response of cancer cells to immune checkpoint inhibition^13,14,17^. Several gene expression-based signatures have been proposed to assess the formation of TLS in primary tumors. A 12-chemokine gene signature for the presence of TLS was initially introduced in colorectal cancers^19^. Subsequently, this signature has been employed in various studies including hepatocellular carcinoma^34,35^, melanoma^36^ and breast cancer^37^. In addition, the Tfh-derived gene signature together with CXCL13 expression has been shown as a TLS marker in breast cancer^38^. Although useful to assess TLS presence in primary tumors, none of these signatures has been derived from transcription profiles of BrM. Regardless of the tumor type and even in the absence of a tumor, immune cells that infiltrate into the brain undergo transcriptional adaptations to the unique microenvironment. Here, we report on a specific gene expression signature that reflects the presence of TLS and (TLS-like) lymphoid aggregates in subsets of BrM derived most commonly from lung cancer and melanoma.

Differential gene expression analysis between the TLS-positive and TLS-negative tumors highlighted the upregulation of functional processes and pathways such as chemokine signaling, lymphotoxin beta signaling and evidence of plasma cell expansion (IGH, IGK genes). The precise mechanism underlying the formation of TLS in metastatic brain niches remains to be fully understood. However, increasing evidence in the literature highlights the involvement of specific signaling pathways, particularly those involving lymphotoxin (LT)-α/β for the generation and maintenance of lymphoid structures. High endothelial venule (HEV) differentiation and formation of organized lymphoid aggregates is mediated through LTB Receptor (LTβR) signaling^39^. LIGHT/TNFSF14, a lymphotoxin-related cytokine, has recently emerged for its role in the promotion of TLS-like aggregates with HEVs *in vivo* via the induction of CCL21 secreted by tumor endothelial cells, thus increasing the influx of T and B cells^40^. In addition, we observed the upregulation of cell adhesion molecules such *ICAM-1* and *VCAM-1* in TLS-positive tumors. Adhesion molecules have been reported to colocalize with PNAd+ HEV and TLS in primary non-small cell lung cancers^41^.

In line with the findings by gene expression-based deconvolution, multiplex immunofluorescence and spatial image analyses revealed lymphoid aggregates containing B and T cells indicative of TLS formation in subsets of BrM tissues. Macrophages were found abundantly within TLS, highlighting the presence of myeloid cell niches in BrMs. Metastasis-associated macrophages exhibit a continuum of macrophage phenotypes, underscoring their complexity and plasticity in BrMs^42^. Similarly, it is crucial to investigate the functional roles of TLS-associated macrophages and their impact on tumor progression.

In conclusion, our results highlight a role of B lymphocytes and formation of TLS in the immune microenvironment of subsets of BrM. In particular among lung cancer BrM, we observed a higher prevalence of B cells and TLS, which appeared to be influenced by previous treatment of the primary tumor and associated with prolonged patient survival after BrM diagnosis. In line with these findings, a recent immunohistochemical analyses of a small cohort (n=17) of patients with BrM from lung cancer reported an association with longer postoperative survival^43^. In addition, another recent study reported on a prognostic role of TLS in patients with breast cancer BrM^45^, although our RNA sequencing and spatial immune cell profiling data indicate only rare TLS formation in breast cancer BrM. Hitherto, the potential role of TLS for the response to immune checkpoint inhibition has not been investigated in BrM patients. It thus remains to be clarified whether the presence of TLS may serve as a putative biomarker for patient stratification to immune therapies, similar to extracranial cancers^16,44^, hence offering promise for promoting precision medicine in BrM treatment.

Our results provide a rationale for deeper exploration of the tumor immune microenvironment in BrM. Moreover, it is imperative to elucidate how distinct molecular subtypes or genetic drivers in cancer cells may differentially influence the extent and cellular composition of the tumor immune microenvironment in BrM from different cancer types. In addition, spatial transcriptomics and proteomics together with B and T cell repertoire analysis may provide deeper mechanistic insights into the modulation of immune responses governed by TLS in BrM.

### Limitations of the study

The data reported in this study are subject to limitations due to the small numbers of tissue samples available from the individual cancer types and the retrospective study approach, including tissue and clinical data collection. Available information on treatment of the investigated patient cohort was sparse, however, pointed towards an influence of previous treatment on the composition of the immune environment in BrM. The limited clinical annotation did not allow to access whether the presence of TLS is associated with response to immune checkpoint inhibition in BrM patients. To circumvent these limitations, studies on larger, uniformly treated and prospectively followed up patient cohorts are needed. For methodological reasons, the comparison of results obtained by bulk RNA sequencing and multiplex immunofluorescence staining in our study was limited by the investigation of spatially distinct areas by each method, which may cause discrepant results due to regional heterogeneity in BrM.

## Acknowledgments

The authors would like to thank B. Friedensdorf, H. Seul and N. Beketow (Düsseldorf) for excellent technical assistance. We thank the Spatial Biology Lab at the Institute of Neurology (Edinger Institute) supported by LOEWE-Center “Frankfurt Cancer Institute”, the German Consortium of Translational Cancer Research (DKTK), Uniscientia-Foundation (Vaduz, Switzerland) and Ludwig Edinger-Foundation (Frankfurt, Germany). The Cancer Registry Northrhine-Westfalia is acknowledged for providing patient outcome data. The study was performed within the framework of the collaborative research project entitled “Preventive strategies against brain metastases” (Coordinator: Prof. Dr. F. Winkler, Heidelberg) that was supported by the priority program translational oncology of the German Cancer Aid (Deutsche Krebshilfe, grant no. 70112507 to YR, DS, IH, HW, AB, KHP, GR) and the Hiege Stiftung - Die Deutsche Hautkrebsstiftung (IH).

## Author contributions

The project was conceived by SSM, YR, BB, KHP, and GR. Methodology was developed by SSM, YR, KHP. Experiments were performed by JF, JM, TS and TTF and supervised by KK, YR, KHP, and GR. Bioinformatical data evaluation was performed by SSM, LM, SS and BB. JF, TF, HW, KL, VM, IH, DS, AB and GR contributed patient samples and data. SSM, JM and YR prepared the figures. SSM and YR wrote the manuscript together with BB, KHP and GR. All authors read and approved the final version of the manuscript.

## Supplementary Tables

**Table S1.** Clinical characteristics of the BrM patient cohort. The table presents clinical information, such as patient’s sex, age at the diagnosis of brain metastasis, localization of the tumor and histological classification.

**Table S2.** Deconvolution of the BrM RNAseq data. The table presents the results from MCPcounter score values for individual cell types for all patients.

**Table S3.** TLS signature score. The table presents the expression values (z transformed) for individual genes comprising the TLS signature.

**Table S4.** Results of mIF analyses from an independent melanoma BrM cohort (n=55).The table lists B (CD20+) and T (CD3+) lymphocyte frequencies and TLS classification.

**Table S5.** Differentially expressed genes in TLS-positive and TLS-negative BrM tumors. Only genes with a foldchange value greater than 0.5 and adjusted p values of less than 0.05 are listed here.

## Supplementary Figures

**Figure S1.**
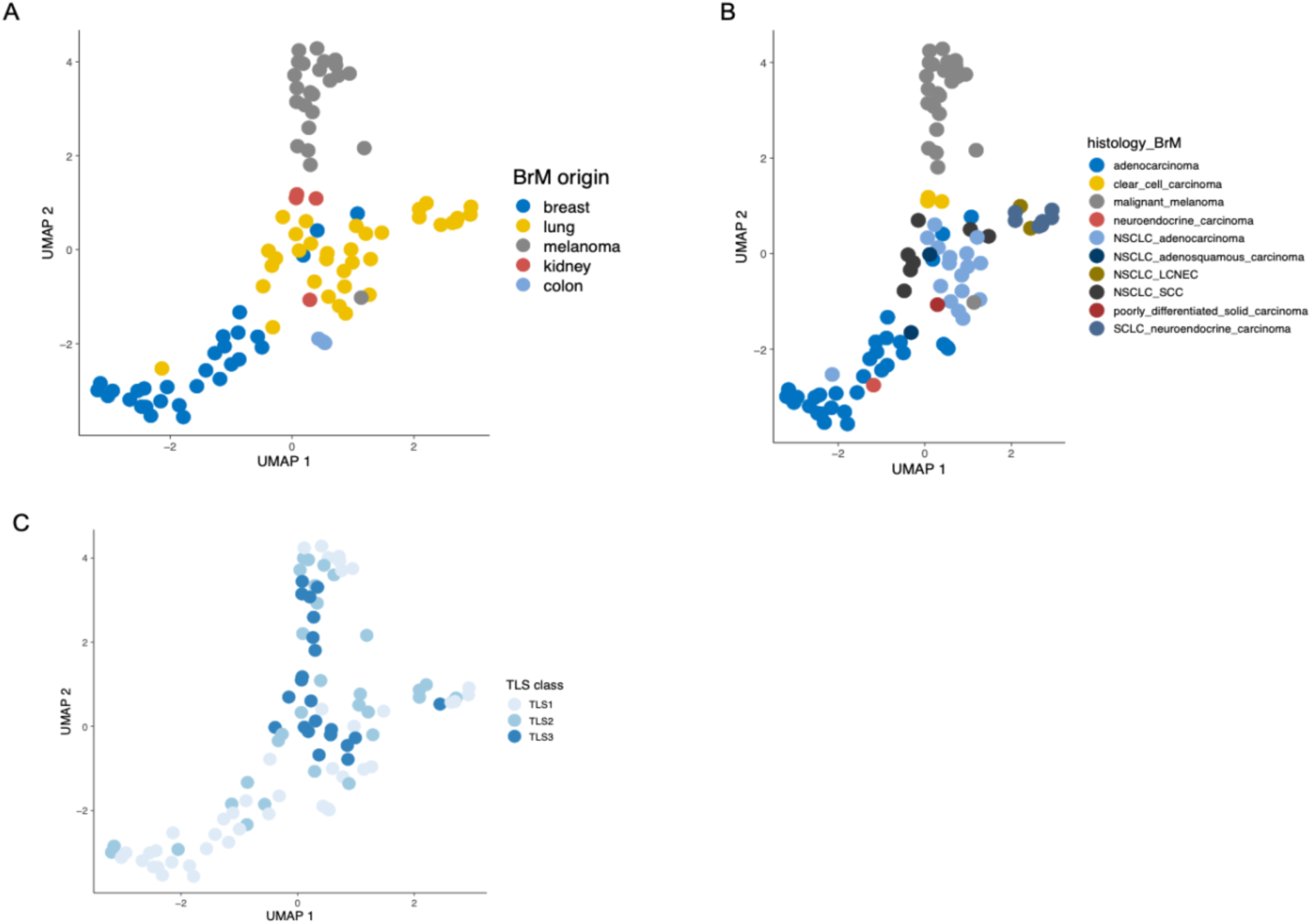
Overview of BrM RNAseq cohort. **(A)** UMAP representation of normalized gene expression data of the BrM cohort according to the primary tumor type of the BrM. **(B)** UMAP representation of normalized gene expression data in the BrM cohort according to the histological classification of the BrM. The lung BrM samples form two clusters and the smaller cluster on the left is mainly composed of BrM from SCLC neuroendocrine tumors and two samples belonging to BrM from NSCLC large cell neuroendocrine carcinoma (LCNEC). **(C)** UMAP representation and assignment of samples to the gene expression-based TLS classes. Color codes for the primary tumors of BrM origin (A), tumor histology (B) and TLS class (C) are provided right to each figure.

**Figure S2.**
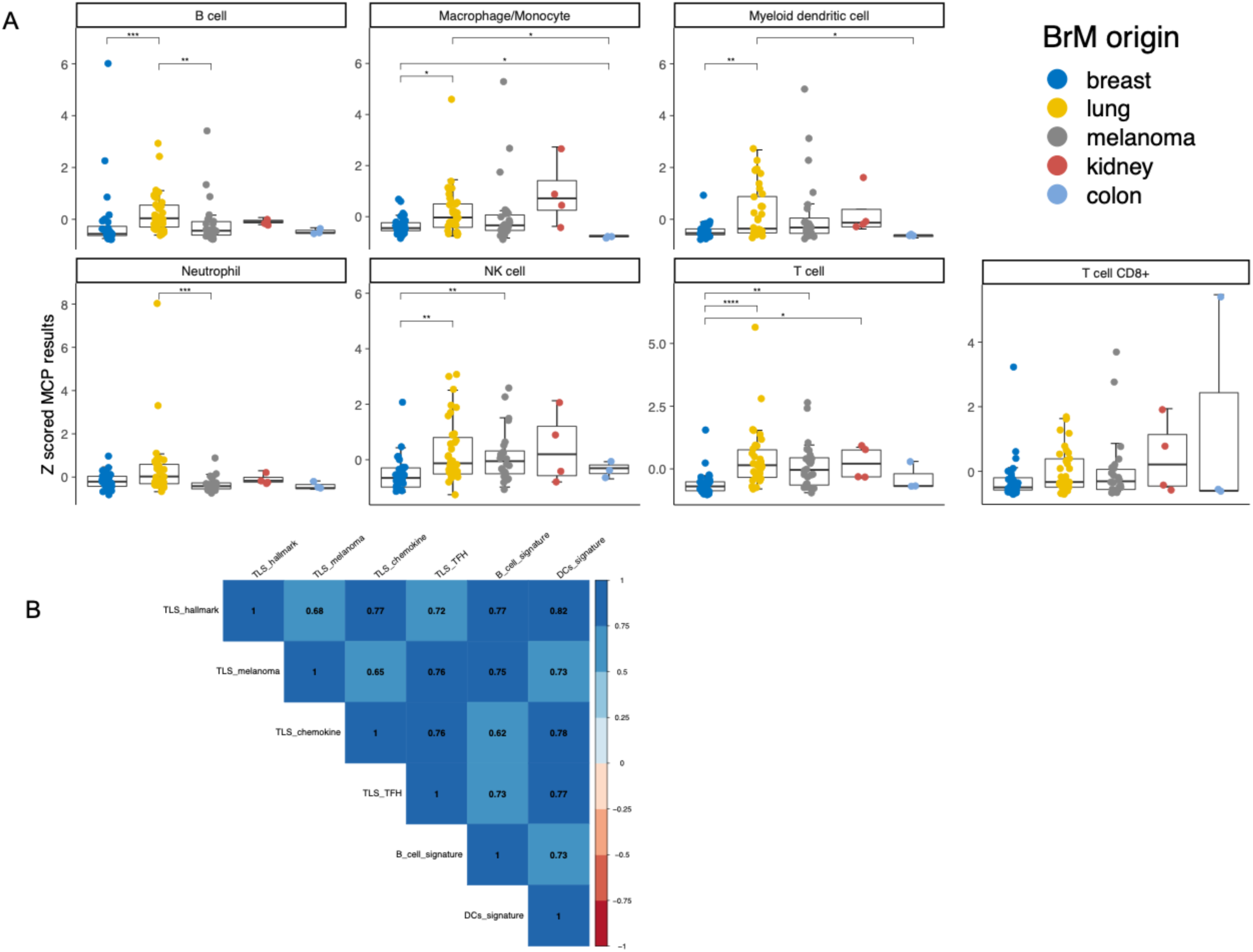
Analysis of tumor infiltrating leukocyte compartments according to cell type using MCPcounter ^18^ immune cell content inference. **(A)** Enrichment of cell states in BrM from different primary tumor types in the investigated BrM cohort of breast, lung, kidney and colon carcinomas as well as melanomas. The p-values were corrected for multiple testing using the Benjamini-Hochberg method. **(B)** Spearman correlation based heatmap depicting the associations between TLS gene signatures in BrM.

**Figure S3.**
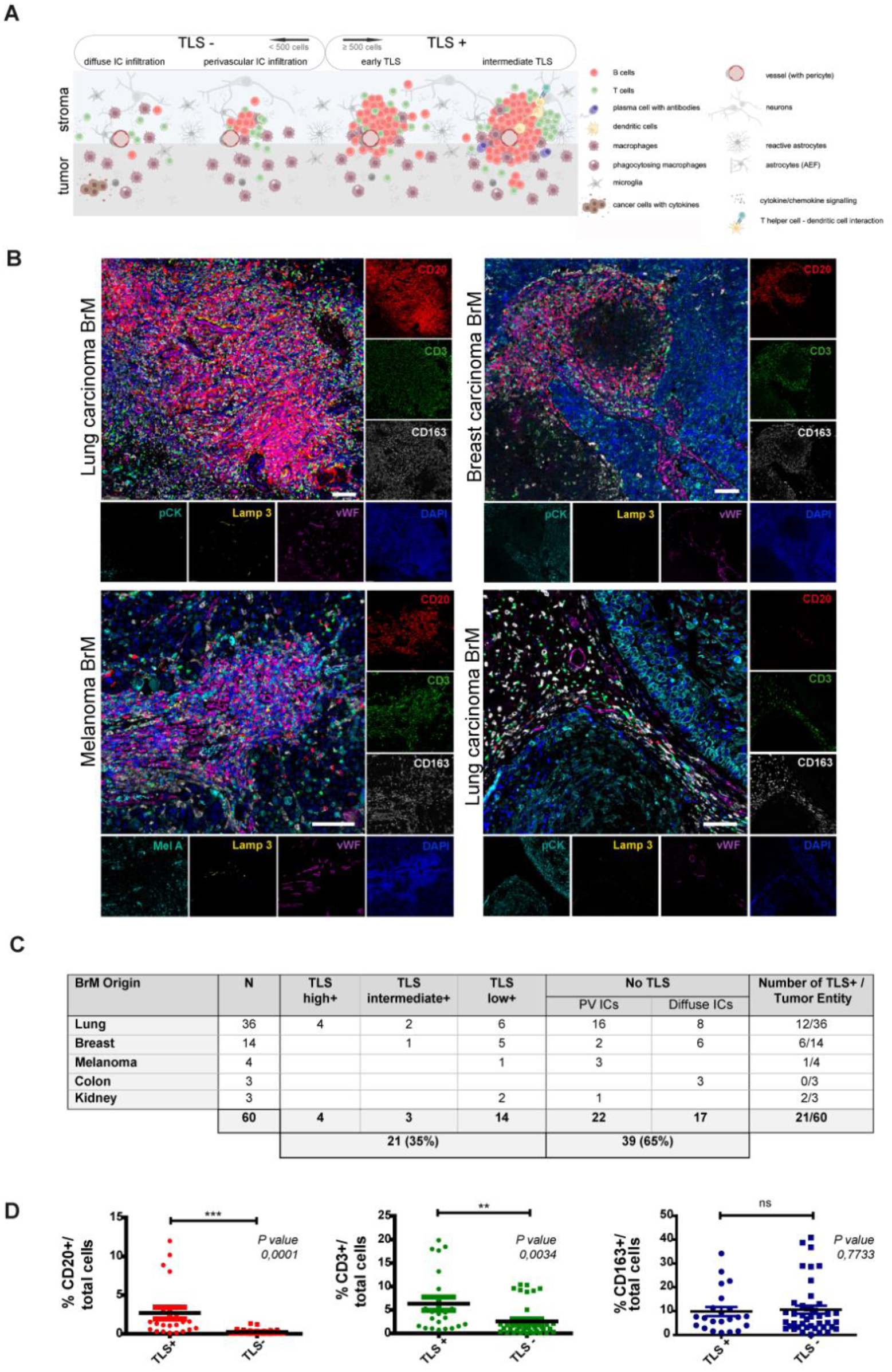
TLS assessment and frequency in BrM of different primary tumors. **(A)** Cartoon displaying immune cell infiltration and aggregation pattern in BrM. TLS were identified as aggregates of CD3+ and CD20+ lymphocytes containing at least 500 cells with a predominance of B cells (>50%; early TLS). In the presence of Lamp3+ dendritic cells, TLS were considered as intermediate TLS. Cases with perivascular immune cell aggregates < 500 cells and diffuse immune cell infiltration were considered as TLS-negative. Cartoon created with BioRender.com. **(B)** Representative composite images of the 7-plex multiplex immunofluorescence (as shown in Fig 3C) with respective monoplex images in selected cases of lung carcinoma BrM, breast carcinoma BrM, and melanoma BrM. Single channel images reveal CD20+ B lymphocytes (red), CD3+ T lymphocytes (green), CD163+ macrophages (white), pCK+ or Mel A+ tumor cells (cyan), Lamp3+ dendritic cells (yellow), vWF+ endothelial cells (magenta) and cell nuclei (blue). Scale bars: 50 µm. **(C)** Table displaying the frequency of TLS in BrM of different primary tumor origins. TLS scores ranged from high over intermediate to low. In TLS-negative cases, small perivascular immune aggregations of T and B cells (<500 cells) and diffuse immune cell infiltration were observed. **(D)** Quantification of CD20+ B lymphocytes, CD3+ T lymphocytes, and CD163+ immunosuppressive macrophages in whole slide multispectral images of BrM (N=60) revealed significantly increased infiltration of B- and T-lymphocytes in TLS+ samples (p=0.001 and p=0.034, respectively) whereas there was no difference in infiltration of immunosuppressive macrophages between TLS+ and TLS-samples (p=0.7733).

**Figure S4.**
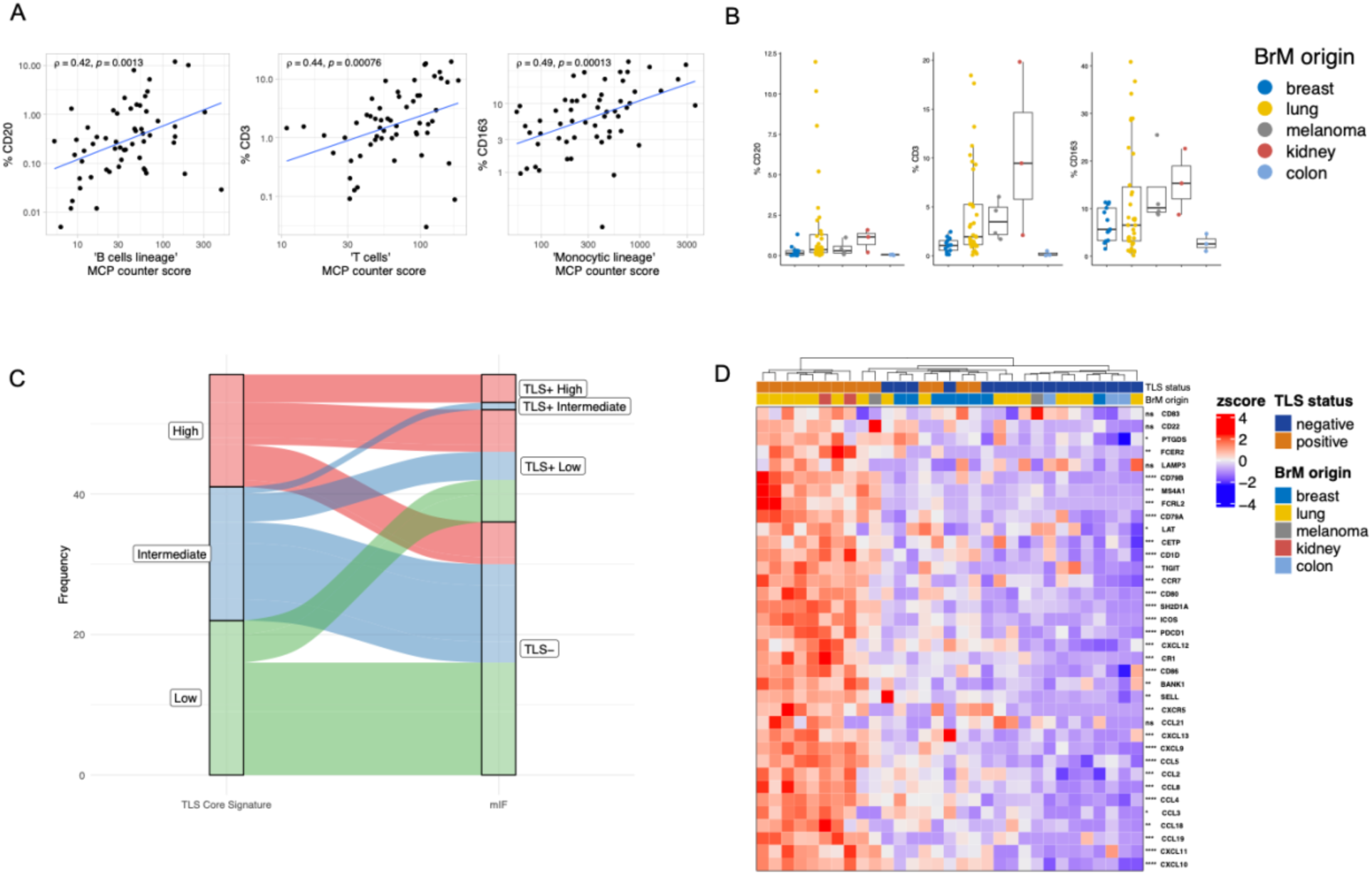
Transcriptomic hints for TLS formation in BrM. **(A)** Correlation of immune cell abundances between the predicted MCP-counter cell scores and multiplex immunofluorescence validations. **(B)** Boxplots displaying the abundances of CD20^+^, CD3^+^ and CD163^+^ cells across BrM from different types by primary cancers. **(C)** River plot showing the overlap of the TLS classification of samples by RNAseq-based TLS signature score and categorization based on multiplex immunofluorescence. **(D)** TLS signature score gene expression in patients with TLS-positive and - negative BrM. The heatmap displays the results of unsupervised hierarchical clustering of the samples (n=31) based on the TLS signature score. Columns and rows are clustered using complete linkage and Euclidean distance.

**Figure S5.**
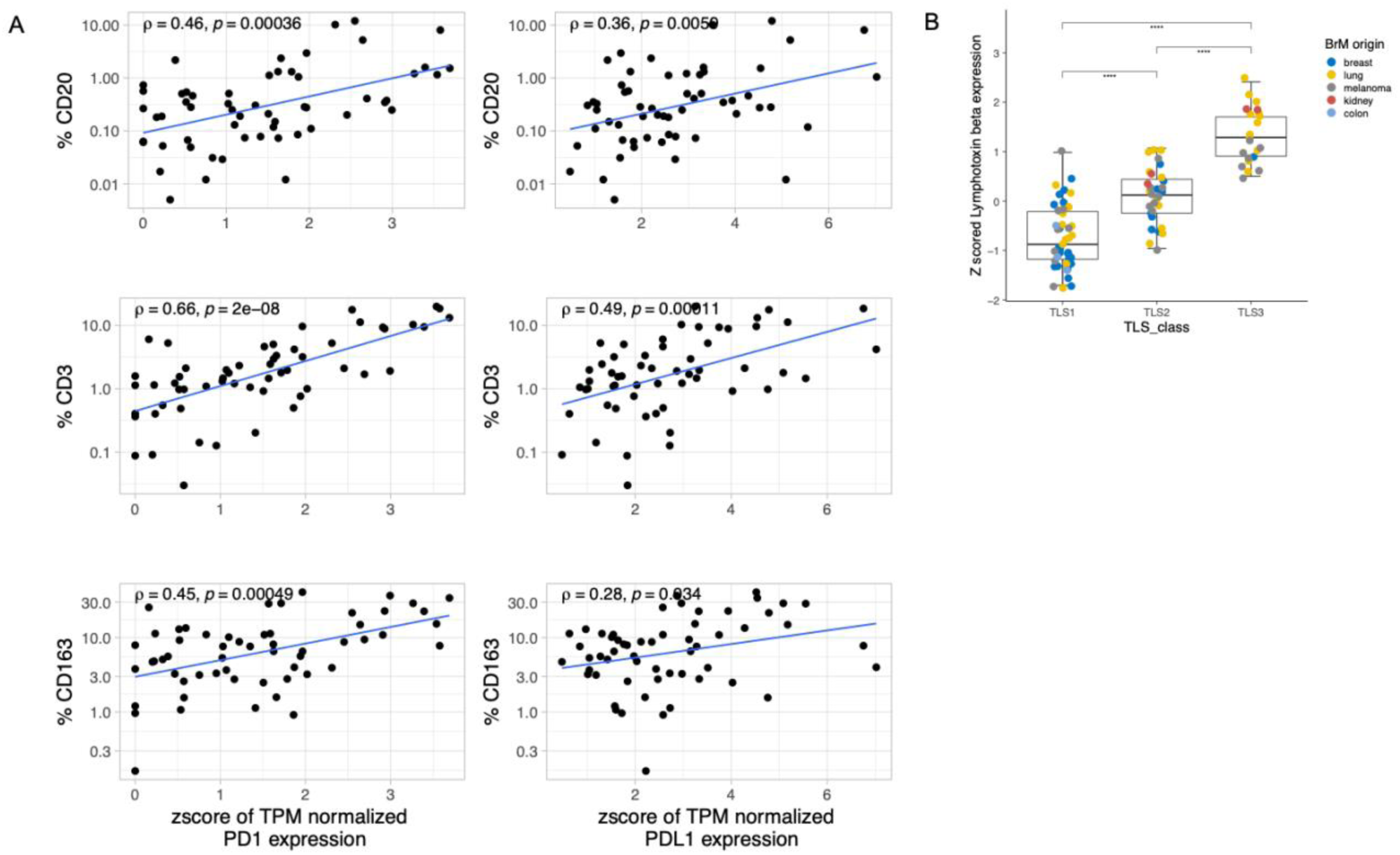
Association between lymphoid cell infiltration and expression of major immunomodulatory genes in BrM. **(A)** Scatter plot displaying the correlation of CD20^+^, CD3^+^ and CD163^+^ cells with PD-L1 and PD-1 gene expression across the investigated BrM samples **(B)** Boxplots displaying normalized gene expression of lymphotoxin beta (LTB) across the TLS classes of BrM. The different BrM origins are highlighted according the color code given right to the graph. Medians are indicated and error bars depict standard deviation. Statistical testing was performed using an unpaired two-sided Wilcoxon test and p-values were corrected for multiple testing using the Benjamini-Hochberg method. Significance levels between groups are: ****, p≤0.0001; ***, p≤0.001; **, p≤0.01; *, p≤0.05; ns (not significant), p>0.05.

**Figure S6.**
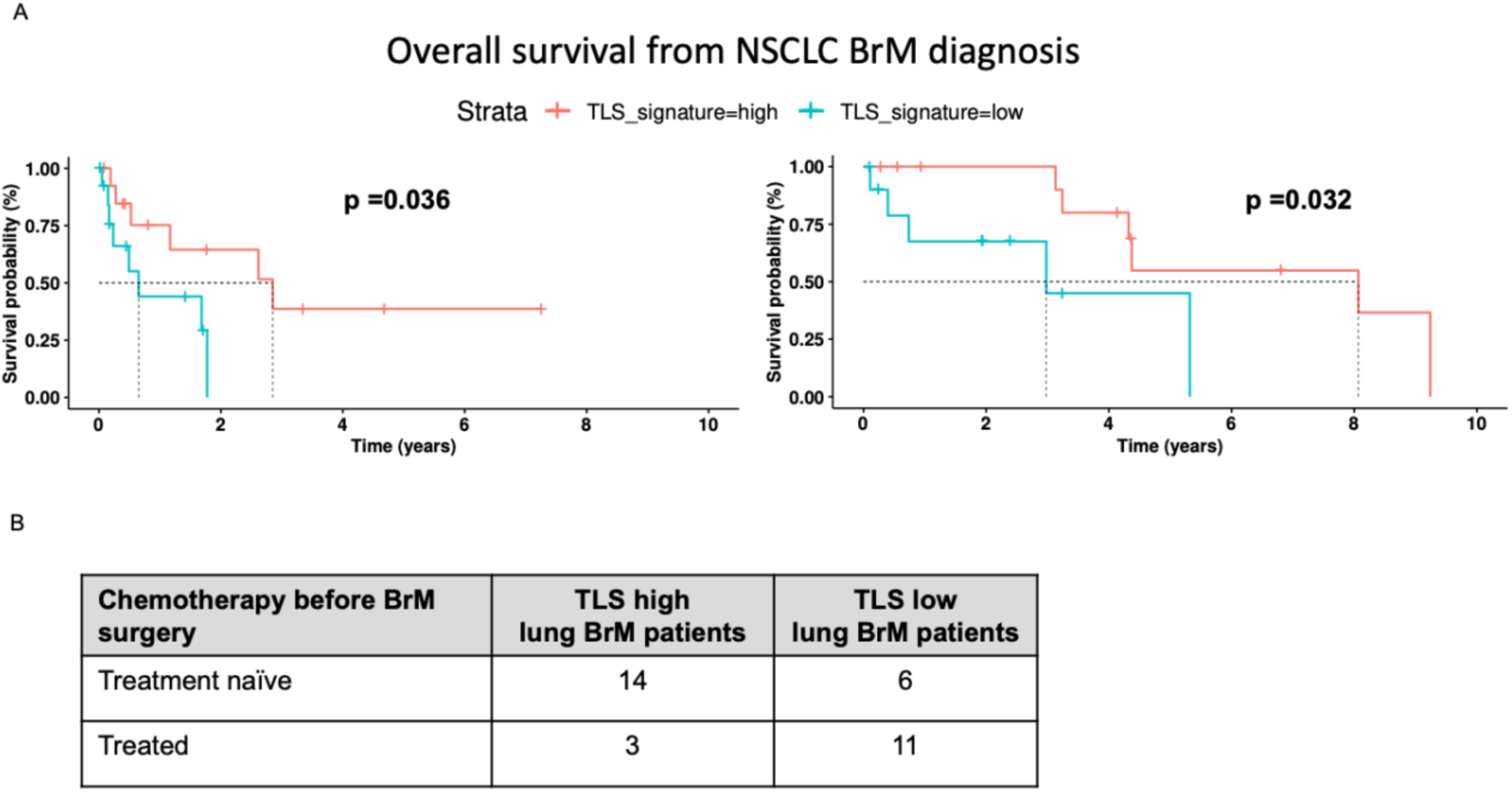
Association between TLS class assignment and previous treatment by chemotherapy in patients with BrM from lung cancer and survival in patients with NSCLC BrM. **(A)** Prolonged survival of patients with BrM from NSCLC with high TLS score in the own institutional cohort (left) and an independent published cohort^24^ (right). **(B)** In the institutional cohort of 34 patients with BrM from lung cancer, 20 patients had treatment-naïve BrM while 14 patients had received chemotherapy before BrM surgery. Logistic regression using a binominal model showed that TLS high group BrM assignment positively associates with treatment-naïve BrM (p=0.00835).

## Extended Methods

### Patients and tissue samples

In total, BrM tissue samples from 95 patients were investigated, including samples from 35 patients with BrM from lung carcinoma (28 patients with NSCLC, 7 patients with SCLC), 29 patients with BrM from breast carcinoma, 24 patients with BrM from cutaneous melanoma, four patients with BrM from renal clear cell carcinoma, and three patients with BrM from colon carcinoma. From each patient, representative tumor tissue samples were fixed in buffered formalin and embedded in paraffin (FFPE) for routine histology and multiplex immunofluorescence analysis. In addition, unfixed tissue specimens were immediately shock-frozen in liquid nitrogen following neurosurgical resection and stored deep-frozen at −80°C until extraction of nucleic acids. Tumors were histologically classified by experienced neuropathologist (GR, JF) based on conventional histological staining (hematoxylin-eosin, PAS, alcian-blue) complemented with immunohistochemical stainings for cytokeratins 7 and 8, TTF1, napsin, synaptophysin, chromogranin A, and MIB1 (lung cancer BrM), cytokeratins 7 and 8, GATA3, estrogen receptor, progesterone receptor, and HER2/neu (breast cancer BrM), melan A, HMB-45, S100 and vimentin (melanoma BrM), cytokeratins 8 and 20, as well as CDX2 (colon cancer BrM), and cytokeratin 8, vimentin, CD10 and PAX8 (renal clear cell carcinoma BrM). Immunohistochemical stainings were routinely performed on an automated immunostainer (Dako autostainer link 48, Agilent Technologies, Santa Clara, CA) using horseradish peroxidase-coupled secondary antibodies and 3.3-diaminobenzidine for visualization of antibody binding as described. The patients gave their written informed consent for the use of their tissue samples and associated clinical data for research purposes. The study was approved by the institutional review board of the Medical Faculty, Heinrich Heine University Düsseldorf (study number: 5717). Table S1 provides an overview of the most important clinicopathological data of the patient cohort. Clinical data, including information on outcome, of the patients with BrM from lung cancer were retrospectively retrieved from the institutional patient files and the Cancer Registry of Northrhine-Westphalia. Among these 35 patients, the primary lung cancer had been diagnosed in 21 patients before BrM surgery and 14 of these patients had received systemic chemotherapy for primary tumor treatment. In 14 patients, the primary lung cancer was detected at the time of BrM surgery or shortly afterward. In 20 patients no systemic treatment had been administered before BrM surgery and none of 34 patients had received immune checkpoint inhibitors before BrM surgery. For one patient, no data on treatment before BrM surgery were available.

An independent study cohort of 53 melanoma patients, 25 women, and 28 men, suffering from malignant melanoma and diagnosed with brain metastases was used to validate the frequency of TLS formation as determined by multiplex immunofluorescence-based immune cell profiling. A detailed description of the patient characteristics has been published before^20^.

For independent validation of the association between TLS signature class and outcome in patients with lung cancer BrM (Figure 4E), we retrieved the expression data set of Rubio-Perez et al.^23^ from the Gene Expression Omnibus (GEO) database under the accession code GSE159407.

### RNA extraction

Deep-frozen unfixed tumor tissue samples used for RNA extraction were histologically evaluated for tumor cell content. Only tumor specimens showing an estimated tumor cell content of ≥ 80% were used for RNA extraction. RNA was extracted using the Maxwell® RSC RNA FFPE kit and the Maxwell® RSC instrument (Promega, Mannheim, Germany) as recommended by the manufacturer.

### RNA sequencing

Total RNA samples used for transcriptome analyses were quantified using the Qubit RNA HS Assay (Thermo Fisher Scientific) and quality was measured by capillary electrophoresis using the Fragment Analyzer and the ‘Total RNA Standard Sensitivity Assay’ (Agilent Technologies, Inc. Santa Clara, USA). A mean RNA quality number (RQN) of 8.2 was calculated across all samples and only samples with a RIN value of ≥ 7.0 were sequenced. Library preparation was performed according to the manufacturer’s protocol using the ‘Illumina® Stranded Total RNA Prep Ligation with Ribo-Zero Plus’. Briefly, 500 ng of total RNA was used as input for rRNA depletion, fragmentation, synthesis of cDNA, adapter ligation and library amplification. Bead purified libraries were normalized and finally sequenced on the NextSeq 2000 system (Illumina Inc. San Diego, USA) with a read setup of PE 2×150 bp. The Illumina DRAGEN FASTQ generation tool (version 3.8.4) was used to convert the bcl files to fastq files, as well for adapter trimming and demultiplexing.

### Alignment of transcriptome sequencing data

RNA sequencing reads were mapped with STAR (version 2.5.3a)^46^.

For building the index, GENECODE version 19 gene model (https://www.gencodegenes.org/human/release_19.html) was used and merging of reads was performed using Sambamba (version 0.6.5). The output was converted to sorted BAM files with SAMtools (version 1.6). The following alignment parameter were used: –twopassMode Basic – twopass1readsN −1 –genomeLoad NoSharedMemory –outSAMtype BAM Unsorted SortedByCoordinate –limitBAMsortRAM 100000000000 –outBAMsortingThreadN=1 – outSAMstrandField intronMotif –outSAMunmapped Within KeepPairs –outFilterMultimapNmax 1 – outFilterMismatchNmax 5 –outFilterMismatchNoverLmax 0.3 –chimSegmentMin 15 –chimScoreMin 1 –chimScoreJunctionNonGTAG 0 –chimJunctionOverhangMin 15 –chimSegmentReadGapMax 3 – alignSJstitchMismatchNmax 5 −1 5 5 –alignIntronMax 1100000 –alignMatesGapMax 1100000 – alignSJDBoverhangMin 3 –alignIntronMin 20.

### Quantification of gene expression

Expression levels were quantified per gene and sample as reads per kilobase (kb) of exon model per million mapped reads (RPKM), and RefSeq (https://www.ncbi.nlm.nih.gov/refseq/) was used as gene model. For each gene, overlapping annotated exons from all transcript variants were merged into non-redundant exon units with a custom Perl script. Nonduplicate reads with mapping quality > 0 were counted for all exon units with coverageBed from the BEDtools package^47^ version 2.16.2. To derive the RPKM value, read counts were summarized per gene and divided by the combined length of its exon units (in kb) and the total number of reads (in millions) as the sum of reads counted by coverageBed.

Size factor and dispersion estimation were calculated for raw count data before performing Wald statistics using DESeq2^48^. Normalized read count values for individual genes were centered and scaled (z-score), and quantile discretization was performed. Complete-linkage analysis with Euclidean distance measure was used for clustering. The heatmaps were generated using the R package ComplexHeatmap^49^. Principal component analysis was performed using singular value decomposition (prcomp) to examine the co-variances between samples. Differential expression between the TLS +ve and TLS-ve samples was performed using DESeq function within DESeq2. Gene set enrichment analysis was performed using the R package clusterProfiler^50^.

### Deconvolution of cell types and tumor microenvironment composition

To estimate the immune cell composition of BrM, we used the MCP-counter tool^18^ to obtain cellular abundance scores for two stromal and six immune cell populations from bulk RNA-sequencing samples. The scores were returned in arbitrary units and hence were comparable across samples.

### Gene signature for tertiary lymphoid structure

We built a metagene signature based on the compendium of TLS-related genes derived from recent studies including the 12-chemokine TLS signature (CCL2, −3, −4, −5, −8, −18, −19, −21, CXCL9, −10, - 11, −13)^51^, B cell markers (BANK1, CD19, CD22, CD79A, CR2, CR1, FCRL2, MS4A1, PAX5, FCER2, MZB1), activated dendritic cell markers including the germinal center light zone (LAMP3, CD80, CD83, CD86, CCR7), TLS signature derived from a melanoma dataset (CD79B, RBP5, EIF1AY, CETP, SKAP1, LAT, CCR6, CD1D, PTGDS)^14^ and TLS-hallmark genes (CCL19, CCL21, CXCL13, CCR7, CXCR5, SELL, LAMP3)^14^ and Tfh signature (CXCL13, CD200, FBLN7, ICOS, SGPP2, SH2D1A, TIGIT, PDCD1)^52^.

To construct the metagene TLS signature for BrM samples, we performed a correlation analysis of each gene with the 12 chemokine TLS signature and selected genes with the highest correlation (Pearson’s correlation). The genes with low expression across all samples or with a negative correlation to the overall signature score were discarded from the final list of genes. Our TLS signature for brain metastases comprised of 32 genes (Table S3). The TLS signature score described above was adapted to analyze published independent samples from nanostring data. A TLS signature score was calculated using the geometric mean of the 32 gene signature. The lung BrM patient cohort was stratified into TLS high score and low score based on the median values among the patient cohort.

### Multiplex immunofluorescence and multispectral image analysis of BrM biopsies

Formalin-fixed and paraffin-embedded (FFPE) sections of BrM specimens were stained by using Opal Polaris multicolour kit (NEL861001KT, Akoya Biosciences, Inc.) based on thyramide signal amplification immunostaining method. Multiplex stainings targeting anti-human CD3 (DAKO, A0452), CD20 (MA516334, Thermo Fischer Scientific), CD163 (Abcam, ab265592), pCK (DAKO, M0821), Lamp-3 (Abcam, ab271053) and vWF (DAKO, A0082) were performed on LabSat™ Research Automated Staining Instrument (Lunaphore Technologies SA). DAPI was used to detect nuclei. Images were acquired on *Vectra Polaris*™ Automated Quantitative Pathology Imaging System (Akoya Biosciences, Inc.) using MOTiF™ technology at 0,5 µm/pixel. Whole slide multispectral image analysis was performed with (I) Phenochart®, version 1.0.12, a whole slide scan viewer, (II) InForm® Image Analysis Software (Akoya Biosciences), which was used for spectral unmixing and batch analysis, and (III) HALO® image analysis software (Indica Labs, Albuquerque, NM), which was used for quantitative spatial analysis of fluorescently unmixed whole slide multispectral images. TLS were identified as perivascular aggregates of CD3+ and CD20+ lymphocytes, containing at least 500 cells with a predominance of B cells (>50%). In the presence of Lamp3+ dendritic cells, TLS were considered more mature and called “intermediate TLS” (adapted from Petitprez et al., 2020)^17^. Cases with perivascular immune cell aggregates < 500 cells and diffuse immune cell infiltration were considered as TLS-negative.

### Statistical analyses

All statistical analyses were performed using R (version 4.0.0) and the packages survival and ggplots. The relationship between a categorical variable and a quantitative variable was estimated with the Mann–Whitney U test (two categories) or the Kruskal–Wallis test (three or more categories). All tests were two-sided. The relationship between two quantitative variables was estimated with the Pearson correlation. P values were corrected for multiple hypothesis testing with the Benjamini–Hochberg method, as indicated in the text and figure legends. Univariate survival analyses were conducted using Kaplan–Meier estimates and log-rank tests.

## Code availability

All codes used in this study are available from the corresponding author upon request.

## Data availability

Sequencing data were deposited in the European Genome-Phenome Archive (EGA) under accession EGAS50000000563. The nanostring data of the Rubio-Perez et al. (2021) BrM cohort^24^ used for Figure 4E and Figure S6A were retrieved from the publicly available Gene Expression Omnibus (GEO) database under accession code GSE159407.

